# A novel combination therapy for ER+ breast cancer suppresses drug resistance via an evolutionary double-bind

**DOI:** 10.1101/2024.09.03.611032

**Authors:** Rena Emond, Jeffrey West, Vince Grolmusz, Patrick Cosgrove, Aritro Nath, Alexander R.A. Anderson, Andrea H. Bild

**Affiliations:** City of Hope, Department of Medical Oncology and Therapeutics Research, Beckman Research Institute, City of Hope National Medical Center, Monrovia, CA, 91016, USA; Integrated Mathematical Oncology Dept. Moffitt Cancer Center, 12902 USF Magnolia Drive, Tampa, FL 33612

**Keywords:** Disulfiram, organoid, chemotherapy resistance, evolutionary game theory, mathematical modeling

## Abstract

Chemotherapy remains a commonly used and important treatment option for metastatic breast cancer. A majority of ER+ metastatic breast cancer patients ultimately develop resistance to chemotherapy, resulting in disease progression. We hypothesized that an “evolutionary double-bind”, where treatment with one drug improves the response to a different agent, would improve the effectiveness and durability of responses to chemotherapy. This approach exploits vulnerabilities in acquired resistance mechanisms. Evolutionary models can be used in refractory cancer to identify alternative treatment strategies that capitalize on acquired vulnerabilities and resistance traits for improved outcomes. To develop and test these models, ER+ breast cancer cell lineages sensitive and resistant to chemotherapy are grown in spheroids with varied initial population frequencies to measure cross-sensitivity and efficacy of chemotherapy and add-on treatments such as disulfiram combination treatment. Different treatment schedules then assessed the best strategy for reducing the selection of resistant populations. We developed and parameterized a game-theoretic mathematical model from this in vitro experimental data, and used it to predict the existence of a double-bind where selection for resistance to chemotherapy induces sensitivity to disulfiram. The model predicts a dose-dependent re-sensitization (a double-bind) to chemotherapy for monotherapy disulfiram.

## Introduction

Identifying two drugs with synergistic action when combined is a promising strategy to combat drug resistance. While combination treatments have shown clinical promise^1^, analytical methods to optimize treatment response remain limited due to heterogeneous response, multi-drug resistance, multiple resistance mechanisms, or simply a lack of available drugs that qualify as synergistic^2,3^. In this work, we implement a drug screening strategy in breast cancer that quantifies the presence of drug synergy in parental and evolved resistance breast cancer cell lines to develop new analysis methods for assessment of combination therapy effectiveness. Importantly, this new analysis method also considers the potential for heterogeneous cell-cell interactions to negate (or promote) any drug synergy that is present. Here, we apply this screening process to identify effective chemotherapy add-on treatments in breast cancer cell lines. The goal of this drug screen is to identify a candidate drug that can be classified as an “evolutionary double-bind,” whereby the mechanism of resistance for a first-line therapy induces a vulnerability to this candidate second-line treatment^4–6^.

Drug combinations can be classified as additive (neutral), synergistic, or antagonistic. Identifying candidate drugs that are collaterally sensitive to first-line treatments represents a promising strategy to mitigate treatment-resistance in antibiotics^7,8^ and cancer^9^. Collateral sensitivity is defined as the increased susceptibility to a particular drug when resistance to a separate drug has increased. Collateral sensitivity is often quantified using comprehensive multi-dose drug response assays, applying a range of monotherapy and combination dosing to characterize the synergy or antagonism between the two drugs using mathematical models^10–12^. These approaches allow for rapid screening to identify drug candidates that may have beneficial efficacy in combination with first-line treatments. However, it is less common to repeat this process for evolved-resistance cell lines to note changes in collateral sensitivity after the onset of resistance. Changes in synergy after resistance or refractory disease remains an open question.

Theoretical models indicate the potential for cell-cell interactions within heterogeneous tumors to either promote or negate drug synergy^13^. While collateral sensitivity assays do provide insight into drug interactions that lead to increased efficacy when in combination, they do not address potential cell-cell interactions. For example, heterogeneous cell populations create the possibility for asymmetric interactions among cells that can alter responses to treatment^14–20^. Cells may interact through 1) competition for resources and space^21^, 2) cooperative production of resources^22^, 3) mutualistic suppression of the immune response^23^, or 4) inhibitory effects on neighboring healthy cells^24^. Thus, here we advocate for direct measurement of cell-cell interactions during the drug screening process to aid in determining the robustness of drug synergy within heterogeneous tumors. To measure cell-cell interactions we employ the Evolutionary Game Assay (EGA)^25^ technique to quantify the frequency-dependent growth rates of cell lines in co-culture under different treatment^19,20,26,27^. Evolutionary game theory has been extensively applied in cancer modeling to quantify cell-cell interactions between sensitive and resistant subpopulations^28,29^, for example, in non-small cell lung cancer ^25^ and breast cancer^30–33^. However, the effect of cell-cell interactions on drug synergy remains an open question.

We hypothesized that an ideal secondary drug to combine with first-line therapy would possess the following characteristics. First, the existence of strong drug-drug synergistic effects. Second, the drug-drug synergy is maintained (or even strengthened) in the presence of evolved-resistance. Third, when co-culturing naive and evolved-resistance lines, any synergistic effects would again be maintained (or strengthened). Finally, the secondary drug would preferentially target evolved-resistance cells while simultaneously leading to total tumor regression. To reach these goals, the process for screening promising drugs must account wholistically for the underlying evolutionary and ecological dynamics at play within a heterogeneous, evolving system.

Once a candidate drug is identified, the question of drug scheduling (e.g. sequential or combination therapy) is addressed using mathematical modeling. Due to the nature of the evolutionary double-bind relying on the resistance mechanism of the first drug, the candidate second-line drug may be highly effective in the context of first-line resistance, but much less effective as a first-line treatment. It remains an open question whether two drugs in an evolutionary double-bind should be given in combination or in sequential therapeutic regimens. Broadly, the development of treatment strategies which attempt to exploit competition between tumor subpopulations as a method of prolonging the emergence of resistance is known as evolutionary therapy^34–38^. Historically, it has been a major challenge to parameterize mathematical models due to lack of experimental characterization of relevant cell lines. Often, this has led to mathematical models relying heavily on untested assumptions about the proliferation cost of resistance, growth dynamics, plasticity, non-cell autonomous interactions, collateral sensitivity, and more. For example, many mathematical models take the assumption that the mechanism of resistance diverts energy and resources away from proliferation, resulting in slower-growing resistant subpopulation. However, in some settings resistant subpopulations may be associated with an increased proliferation^39^. Similarly, mathematical models often implicitly assume either neutral or no competition between drug-sensitive and drug-resistant subpopulations. Thus, it is appropriate to design a drug screening process that quantifies these ecological and evolutionary dynamics under treatment.

### Screening for candidate evolutionary double-bind drugs in ER+ breast cancer

Breast cancer is a leading cancer cause of death worldwide. Breast cancer is classified into three main subtypes with hormone receptor-positive tumors (including both estrogen and progesterone receptors) consisting of 75% of these cases^40^. Roughly 20-40% of ER+ patients eventually develop distant metastases and account for the majority of metastatic cancer cases^41^. First line treatment for ER+ breast cancer is endocrine therapy. However, approximately 90% of all patients with MBC eventually develop resistance to endocrine therapy^42,43^. Chemotherapy is a treatment for advanced breast cancer that targets cell proliferation, and is considered a critical drug class in the metastatic setting; therefore, identifying strategies to optimize chemo-effectiveness is critical.

From a panel of several drug candidates, disulfiram is identified as a promising monotherapy to further test in combination with chemotherapy and assess potential treatment synergy. We hypothesize that disulfiram will aid in maintaining treatment sensitivity and allow for prolonged response for follow up chemotherapy cycles. This study uses an in vitro 3D coculture model and evolutionary game theoretic model to optimize treatment regimens in ER+ breast cancer cell spheroids consisting of resistant and sensitive cells, taking advantage of evolutionary dynamics and collateral sensitivity.

We begin by screening for promising candidate drugs (**figure 1**), then quantify synergy of the most promising candidate, disulfiram (**figure 2, table 1**). Next, we perform an evolutionary game assay to quantify cell-cell competition across treatment conditions. An integrative mathematical-experimental analysis predicts that a high dose combination of both drugs (chemotherapy and disulfiram) in tandem maximally suppresses both resistant and treatment-naive cells. Finally, we validate the approach by considering alternating (sequential) therapy strategies, which underperform combination treatment, as predicted. In our assessment of disulfiram as a candidate combination therapy with chemotherapy, we classified it as an evolutionary double-bind due to metrics of 1) drug-drug synergy and 2) sensitive-resistant cell-cell interactions.

**Table 1:**
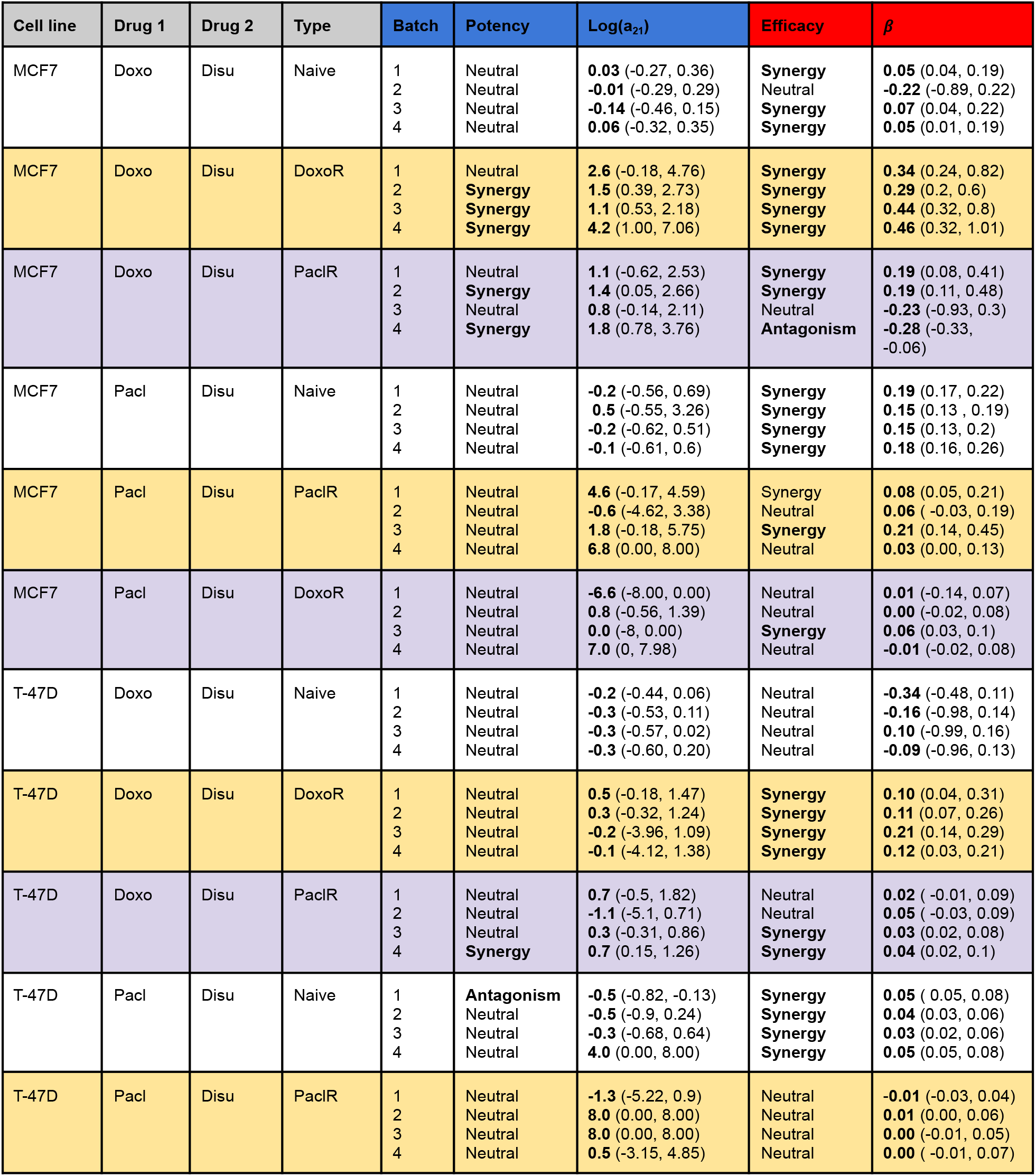
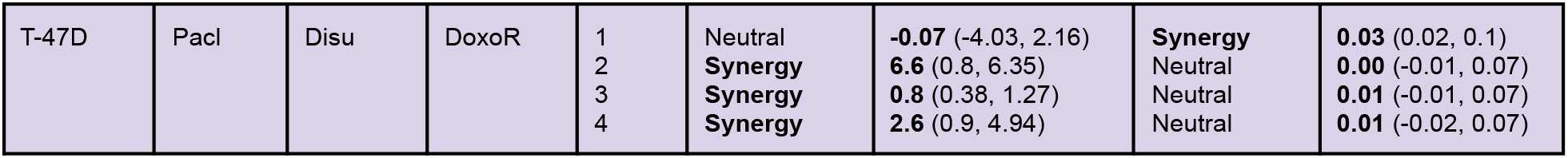
Quantification of drug synergy for two cell lines (MCF7,T-47D) when confronted to two drugs: chemotherapy (paclitaxel or doxorubicin) in combination with disulfiram. Cell lines are either naive, or evolved resistance to paclitaxel or doxorubicin. Each dose response checkerboard assay is repeated four times (batch numbers 1-4 are related to different concentrations of disulfiram, in increasing concentrations) and synergy is measured using the MUSYC software package^11,12^. We consider the effect of disulfiram on the EC50 (potency; log(a_21_): see Methods) and on maximal response (efficacy; *β*: see Methods)

**Figure 1:**
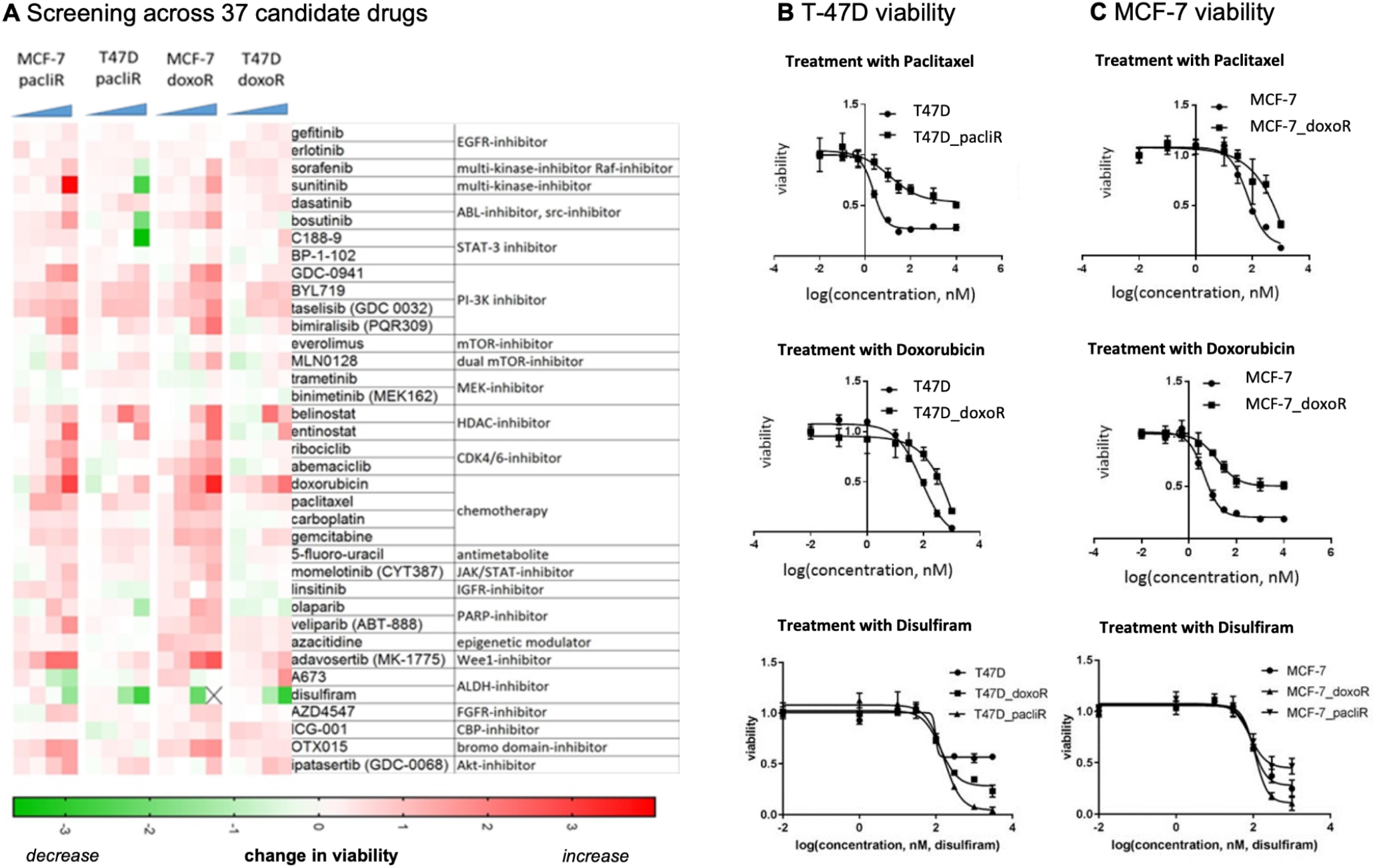
Candidate drugs screened for potential synergistic combinations. **A**, Drug screen performed on MCF7 and T47D chemo resistant cell lines. 37 candidate drugs of different classes of inhibitors, modulators, and chemotherapy were used across a range of treatment concentrations. The decrease in relative cell viability, normalized to the respective maternal sensitive cell line, is shown (green indicating highest decrease/effect in viability). **B**, Growth curves of T-47D sensitive and chemo-resistant cell populations treated with monotherapy paclitaxel, doxorubicin, or disulfiram at various concentrations. **C**, investigation repeated for MCF-7 cell lines.

**Figure 2:**
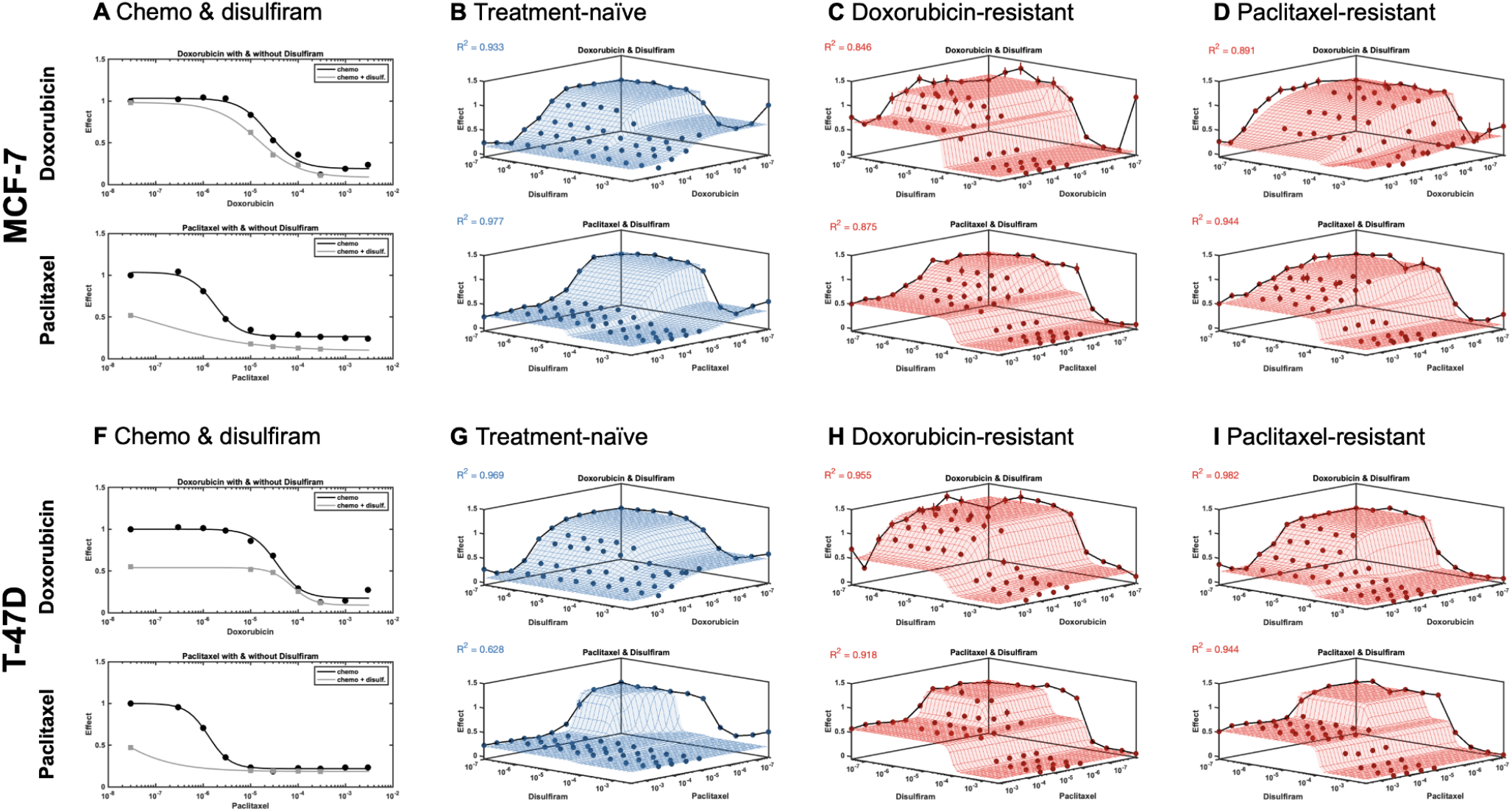
Dose response for MCF-7 and T-47D lines confronted to chemotherapy (doxorubicin and paclitaxel) in combination with disulfiram. **A**, MCF-7 dose response for chemo alone (black) and combination with disulfiram (gray). **B**, Synergy is calculated using MUSYC dose response framework ^11^ for treatment-naive line. repeated for **C**, doxorubicin evolved-resistant line and **D**, paclitaxel evolved-resistant line. **E-H**, Disulfiram investigation repeated for T-47D. Synergistic potency (a_12_, a_21_) and efficacy (*β*) are shown in **Table 1**.

## Results

### Candidate drugs screened for potential synergistic combinations

Thirty-seven candidate drugs were screened to determine the potential for synergistic combinations with chemotherapy across two cell lines (MCF-7, T-47D) with evolved resistance to two different chemotherapy drugs (paclitaxel and doxorubicin). The heatmap in **figure 1A**, indicates potential collaterally sensitivity drugs (green boxes) which improve response in resistant lines. The panel of drugs chosen included several therapies, either FDA approved for oncological use or tested in clinical trials. These include known targeted therapies such as EGFR, PIK3CA/AKT/mTOR, HDAC, CDK 4/6 inhibitors (among others), chemotherapy, and other drugs showing promising off-label use in various oncological settings^44,45^. Candidates with strong drug effect on chemo-resistant cell lines, represented by a decrease in relative cell viability normalized to the maternal sensitive cell line, are shown in green. Disulfiram (an ALDH inhibitor) is a clear outlier for potential synergy across chemo-resistant lines. Follow-up dose-response experiments in sensitive and doxorubicin- and paclitaxel-resistant cell lines confirmed both resistance to mono-chemotherapy as well as the acquired sensitivity to disulfiram in resistant lines compared to the parental sensitive line (**figure 1B, 1C**). In addition, chemo-resistant cells exhibited decreased viability when confronted with disulfiram treatment in a concentration dependent manner. While this was observed in both T47D and MCF7 chemo-resistant cell lines, there were differences in sensitivity to disulfiram between the specific chemo-resistant lines. For example, in T47D, paclitaxel resistant cells harbored a greater sensitivity to disulfiram compared to doxorubicin resistant cells (**figure 1B**). Whereas, MCF7 doxorubicin resistant cells were more negatively affected by disulfiram compared to the paclitaxel resistant line (**figure 1C**). This suggests the molecular resistance mechanisms arising from different chemo-resistant specific phenotypes influence the degree of disulfiram’s targeted drug effect. The range of drug response between the cell lines highlights the importance of concentration considerations during treatment and across different resistant profiles. These findings from our comprehensive drug screen suggest disulfiram as a favorable lead candidate for potential synergistic combination treatment with chemotherapy.

### Quantifying collateral sensitivity

After disulfiram was shown to improve response in chemo-resistant lines, potential synergistic combination treatment with either paclitaxel or doxorubicin was next explored. T-47D and MCF-7 sensitive and resistant lines were confronted with a combination treatment of disulfiram with chemotherapy at various monotherapy and combination concentrations, and the drug effect was assessed by a measure of decreased cell viability. An observed increase in efficacy was seen in both lines when combining chemotherapy with disulfiram treatment compared to mono-chemotherapy (**figure 2A, 2F**). However, with a simple one-dimensional dose response curve, it is not known whether this increase is strictly additive, or if there exists synergistic or antagonistic effects between the two drugs.

To assess and quantify synergy (or antagonism), we performed dose response assays for MCF-7 and T-47D sensitive lines, treated with either doxorubicin or paclitaxel and disulfiram (**Figure 2B, 2F**) as well as MCF-7 and T47D chemo-resistant lines, treated with their corresponding chemo of resistance and disulfiram (**Figure 2C, 2D, 2G, 2H**). This data was subsequently fed into the Multi-dimensional Synergy of Combinations (MUSYC) framework^11^. MUSYC quantifies two types of synergy: synergistic potency (fold-change in the EC-50 half-max concentration when adding the drug) and synergistic efficacy (percent decrease in max effect at high concentrations of both drugs)^10,12^. In treatment-naive lines, the chemo-disulfiram combination is typically neutral (neither synergistic nor antagonistic) in efficacy and potency, suggesting an additive effect that exists between the drugs. This neutrality remains across most evolved-resistant cell lines, with the exception of doxorubicin-resistant MCF-7 cells (**figure 2C**) which shows synergistic potency between doxorubicin and disulfiram. A full description of drug synergy is shown in **Table 1**.

This analysis indicates that disulfiram is an effective drug in combination with chemotherapy strictly due to additive (or at best, weakly synergistic) effects. Importantly, we have confirmed that additivity is maintained even after evolved resistance (**figure 2C,D,G,H**). It’s also important to note that combination (high dose of both chemotherapy and disulfiram) results in maximum efficacy across all cell lines, chemotherapies, and resistance settings.

Interestingly, chemotherapy has an additional stabilizing effect on disulfiram. Visually, disulfiram has a bifurcated effect at high doses when given in isolation (e.g. high doses of disulfiram alone in **figure 2B,C,D,F**), but stabilization of efficacy is achieved when disulfiram is given in combination with chemotherapy. Since the MUSYC framework assumes a non-decreasing effect, it cannot account for this stabilization effect. This analysis ignores any cell-cell interactions between chemo-resistant lines and treatment-naive lines, which we consider in the next section.

### Quantifying cell-cell interactions

Thus far, we have considered treatment-naive cell lines and resistant cell lines in isolation. Next, we perform the Evolutionary Game Assay^25^ as a method for quantifying the fitness of cell lines in co-culture and under different therapies. Here, we define fitness to be synonymous with the growth rate of a cell line. We quantify changes in growth rate as a function of the frequency of competing cell types, known as frequency-dependent growth dynamics.

By labeling the resistant and sensitive cell lines with different fluorescent proteins, this *in vitro* model allowed for the analysis of each cell lines’ different proliferative dynamics and competition in a 3D coculture system using imaging and fluorescence intensity tracking relative to growth. The 3D cell culture allows cells to create cell-cell interactions which resembles more closely the in vivo ecosystem, therefore results obtained from 3D cell culture are clinically more relevant^46^. Cells are seeded across a range of initial resistant (chemotherapy-resistant, R) to naive (treatment-sensitive cells, S) ratios: 0%R, 25%R, 50%R, 75%R, and 100%R. The exponential growth rate is measured for each initial condition, and repeated for untreated control, doxorubicin, disulfiram, and combination doxorubicin and disulfiram (**Fig 3A, C, E, G** respectively). The collection of experiments gives the cell fitness (e.g. growth rate) as a function of the fraction of resistant cells (**Fig 3B, D, F, H**).

**Figure 3:**
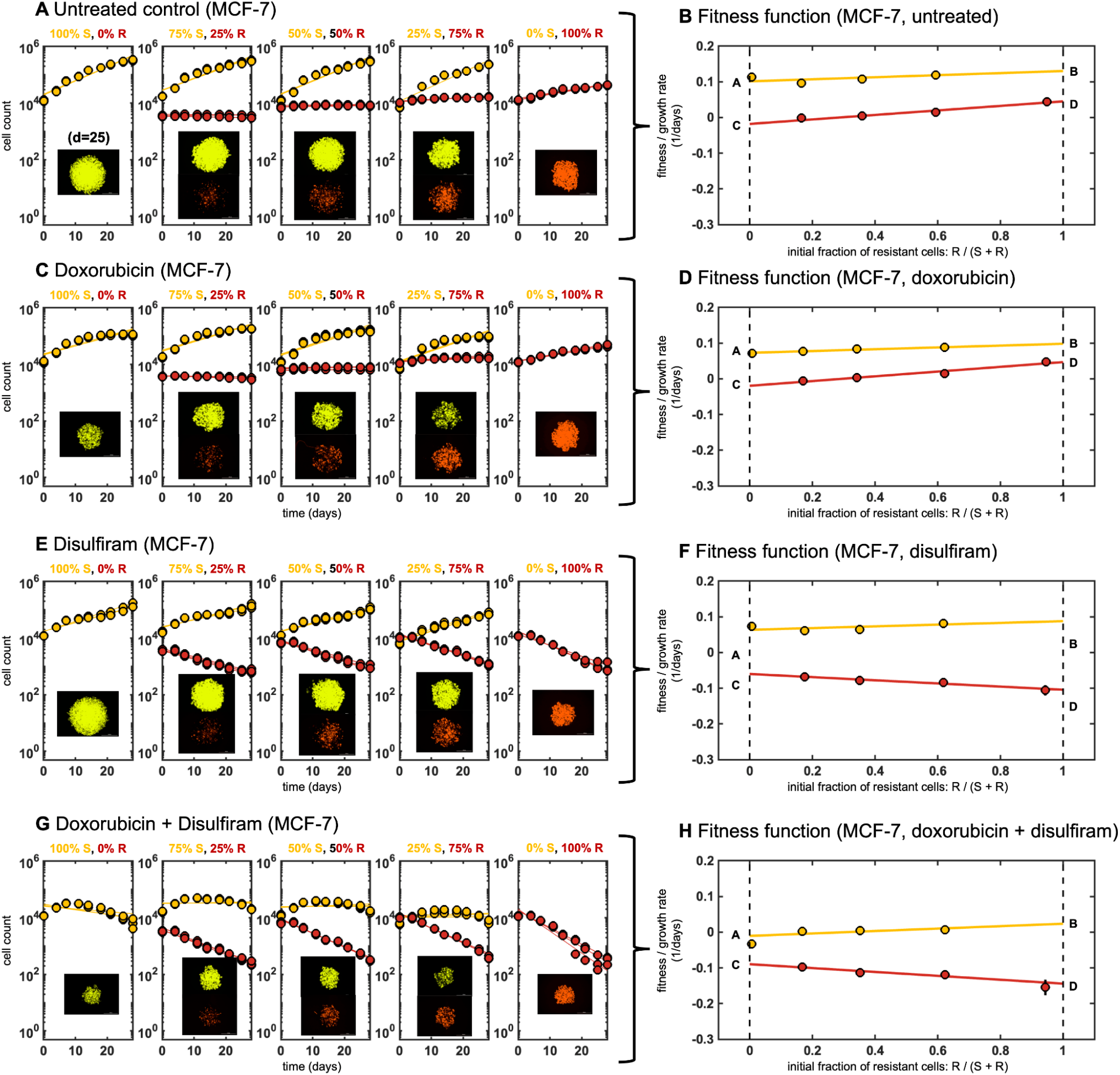
Evolutionary game assay: Cells are seeded with ratio of resistant to naive of 0%R, 25%R, 50%R, 75%R, and 100%R. **A**, The best fit exponential growth rate is measured for each ratio with no treatment. **B**, These growth rates determine the cell type’s fitness as a function of resistant fraction. A linear fitness function is fit with parameters A, B, C, D denoted (See Methods). The process is repeated for spheroids treated with **C-D**, doxorubicin, **E-F**, disulfiram and **G-H**, doxorubicin and disulfiram combination therapy.

In untreated conditions, naive cells tend to grow faster across all initial conditions, suggesting a cost (i.e. a reduction in growth rate) to the mechanism of resistance to chemotherapy (**Fig 3A**). Reduced resistance growth is seen across cell lines (MCF-7, T-47D) and across chemotherapies (doxorubicin, paclitaxel), see Supplementary Data for more details. Naive cells maintain a higher fitness in untreated conditions, a result of a higher proliferation rate, and a competitive advantage (**Fig 3B**). Somewhat unsurprisingly, chemotherapy decreases the fitness of naive cells to a greater extent than resistant cells (**Fig 3C**), across all initial resistant fractions (**Fig 3D**). As expected, chemo-resistant cells are better adept to withstand the effects of chemotherapy compared to sensitive cells. In contrast, the introduction of disulfiram decreases the fitness of resistant cells to a greater extent (**Fig 3E**), across all fractions (**Fig 3F**). When given in combination, chemotherapy and disulfiram together negatively impact the fitness of both populations in coculture (**Fig 3G**), and the additive effect of both drugs decreases the overall total cell count (**Fig 3H**).

Growth dynamics can be classified as having frequency-dependence if the fitness function changes with the seeded ratio (frequency) of resistant cells. Data for each coculture ratio is fit using a linear function (solid red, yellow lines in **Fig. 3B,D,F,H**). The fitness functions shown in **Figure 3** are characterized by two parameters for each cell line (four total parameters): growth rate for monoculture 100% sensitive and 100% resistant (dashed lines in **Fig. 3B,D,F,H**). Fitness (growth rate) is a linear function between the monoculture growth rates for sensitive (labeled A, B) and resistant lines (labeled C, D). These four parameters comprise the payoff matrix (see Methods), providing a complete characterization of the fitness function of two competing cell lines. Thus, we define two important quantities that determine the selection dynamics in this model system: Δ_*S*_ and Δ_*R*_.

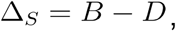

and

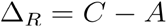

These two quantities represent the invasion fitness for each cell type S, R when the prevalence of that cell type is small. For example, if Δ_*R*_ > 0 resistant cells are expected to invade a population of sensitive cells. Whether that invasion continues to full dominance (resistant cells fully take over) depends on the value of Δ_*S*_ (see Methods).

### Classifying selection dynamics

Figure 4. quantifies Δ_*S*_ and Δ_*R*_ for a range of mono and combination therapy options for MCF-7 with doxorubicin-resistance, paclitaxel-resistance (**Fig 4A, B**) and T-47D with paclitaxel-resistance (**Fig 4C**). MCF-7 sensitive relative fitness (normalized growth rate) is always positive while resistant relative fitness (normalized growth rate) is always negative (Δ_*S*_ > 0, Δ_*R*_ < 0). Disulfiram can be classified as a chemo-sensitizer, due to an increase in sensitivity with increasing disulfiram dose for both monotherapy and combination with chemo (**Fig 4A, B**, left panel). This result is summarized in the middle panels of **Fig 4A** and **Fig 4B**, showing the four possible scenarios for selection dynamics: selection for sensitive (yellow quadrant), co-existence (orange), bistability (blue), or selection for resistance. In MCF7 lines, selection for sensitive cells is always observed under all treatment conditions in coculture, but introduction of disulfiram further shifts and strengthens the selection dynamics in favor of sensitive populations (**Fig 4A, B**, middle panel). Selection for resistance is seen in the T47-D line with chemotherapy in monotherapy or combination (**Fig 4C**).

Next, we introduce a metric for the overall tumor fitness, *ϕ*\*, defined as the long-term growth rate once the selection dynamics reach equilibrium (see **Methods**). Despite defining the selection dynamics, the overall spheroids may be growing (positive tumor fitness, *ϕ*\* > 0) or decaying (negative tumor fitness, *ϕ*\* > 0). For example in all three cell lines, the untreated conditions select for sensitive lines, but maintain positive tumor fitness. In contrast, combination treatment of chemo with maximum disulfiram concentration leads to the lowest (negative) tumor fitness, indicating a negative tumor growth. Note: en route to tumor regression, the model predicts MCF-7 lines become more sensitive (Δ_*S*_ > 0, Δ_*R*_ < 0) while T-47D lines become more resistant (Δ_*S*_ < 0, Δ_*R*_ > 0).

**Figure 4:**
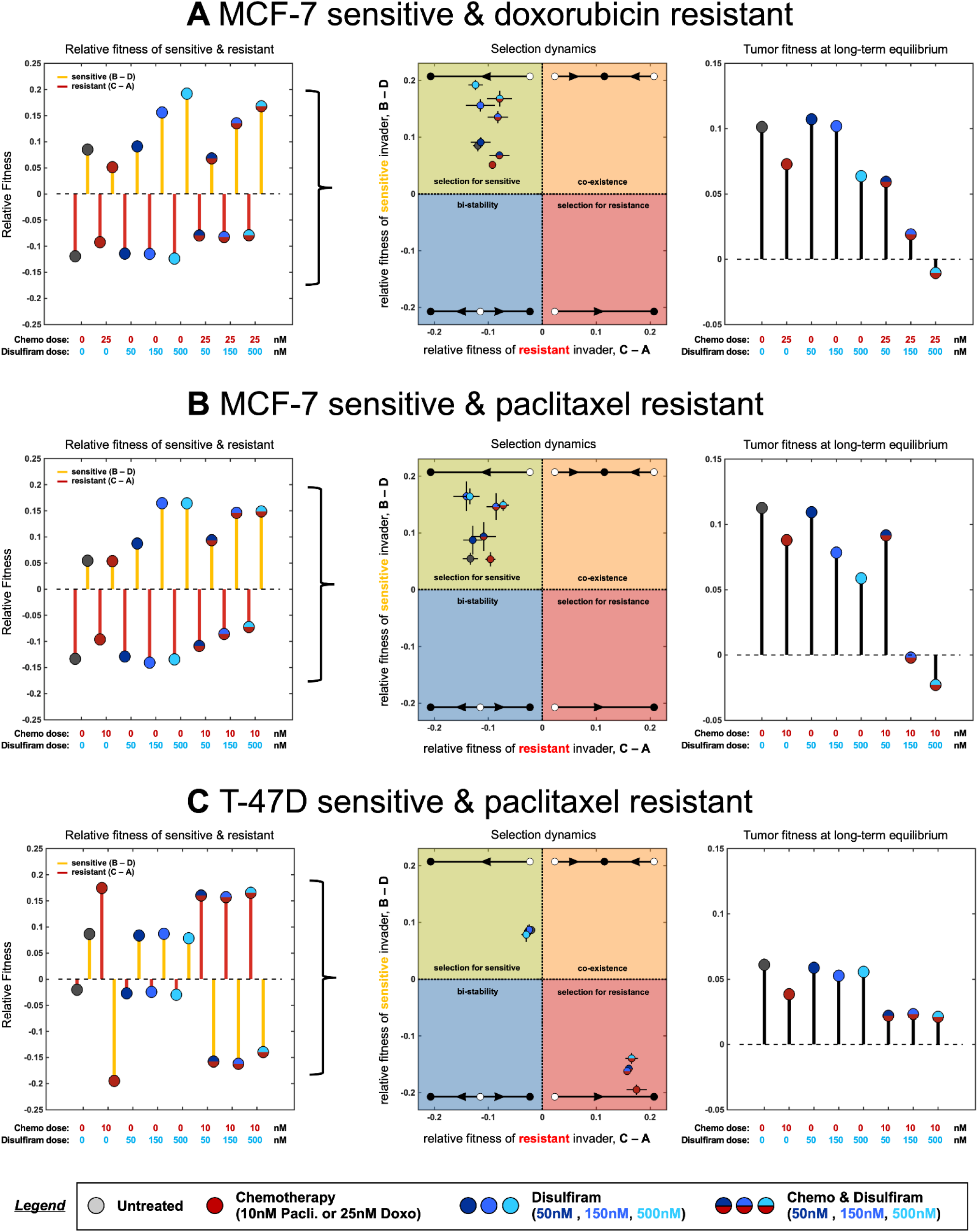
Fitness parameters determine selection dynamics: **A**, MCF7 sensitive and doxorubicin-resistant lines, Left (relative fitness): fitness functions fit in figure 3, shown across all monotherapy or combination therapy options. Relative fitness of sensitive (parameters B - D) and resistant (parameters C - A) shown. Middle (selection dynamics): relative fitness plotted in a quadrant showing selection dynamics as a function of B - D and C- A. All treatment scenarios select for sensitive to varying degrees. Right (tumor fitness): overall tumor fitness of mixed populations shown (long-term model prediction; see Methods). High dose combination therapy minimizes tumor fitness. **B**, relative fitness, selection dynamics, and tumor fitness are shown for MCF7 sensitive and paclitaxel-resistant cell lines. **C**, relative fitness, selection dynamics, and tumor fitness are shown for T47D sensitive and paclitaxel-resistant cell lines.

### Evolutionary Game Assay Validation

Thus far, we have considered only continuous therapeutic regimens: continuous dosing of a single therapy (chemotherapy or disulfiram alone) or continuous dosing of combination chemotherapy with disulfiram. As the mathematical model underlying the EGA technique is parameterized to continuous dosing data, it remains an open question if it’s useful to identify promising sequential therapeutic options (e.g. chemotherapy followed by disulfiram). Next, we consider a range of candidate sequential treatment regimens: alternating chemotherapy and disulfiram with fast (every week) or slow (every two weeks) drug switching. We do not consider any sequential schedule with disulfiram first, as previous figures indicate disulfiram performs best after chemo-resistance is selected for. Note: results from figure 4 intuitively suggest that sequential schedules are unlikely to outperform combination treatments, because tumor fitness (growth rate) is maximally negative under combination treatment, but positive under monotherapy of either chemo or disulfiram (**figure 4A, B, C**; right panels).

This intuition is confirmed in mathematical model simulations in **figure 5**, where combination treatment outperforms these sequential therapeutic regimens across parameterizations for each cell line (**figure 5A, B, C**; top panel). To explore the validity of these *in silico* predictions, our 3D cell culture model was confronted with the different drug treatments. Fluorescently labeled sensitive and chemo-resistant cells were once again plated at various population frequencies previously outlined in figure 3. Co-cultured spheroids were then treated with 25nM of doxorubicin and/or 500nM of disulfiram in continuous combination therapy or alternating/sequential therapeutic strategies. Each sequential regimen explored the possibility of leading treatment with either disulfiram or chemotherapy in an effort to investigate early selection trends of resistant and sensitive populations dependent on initial drug pressure.

**Figure 5:**
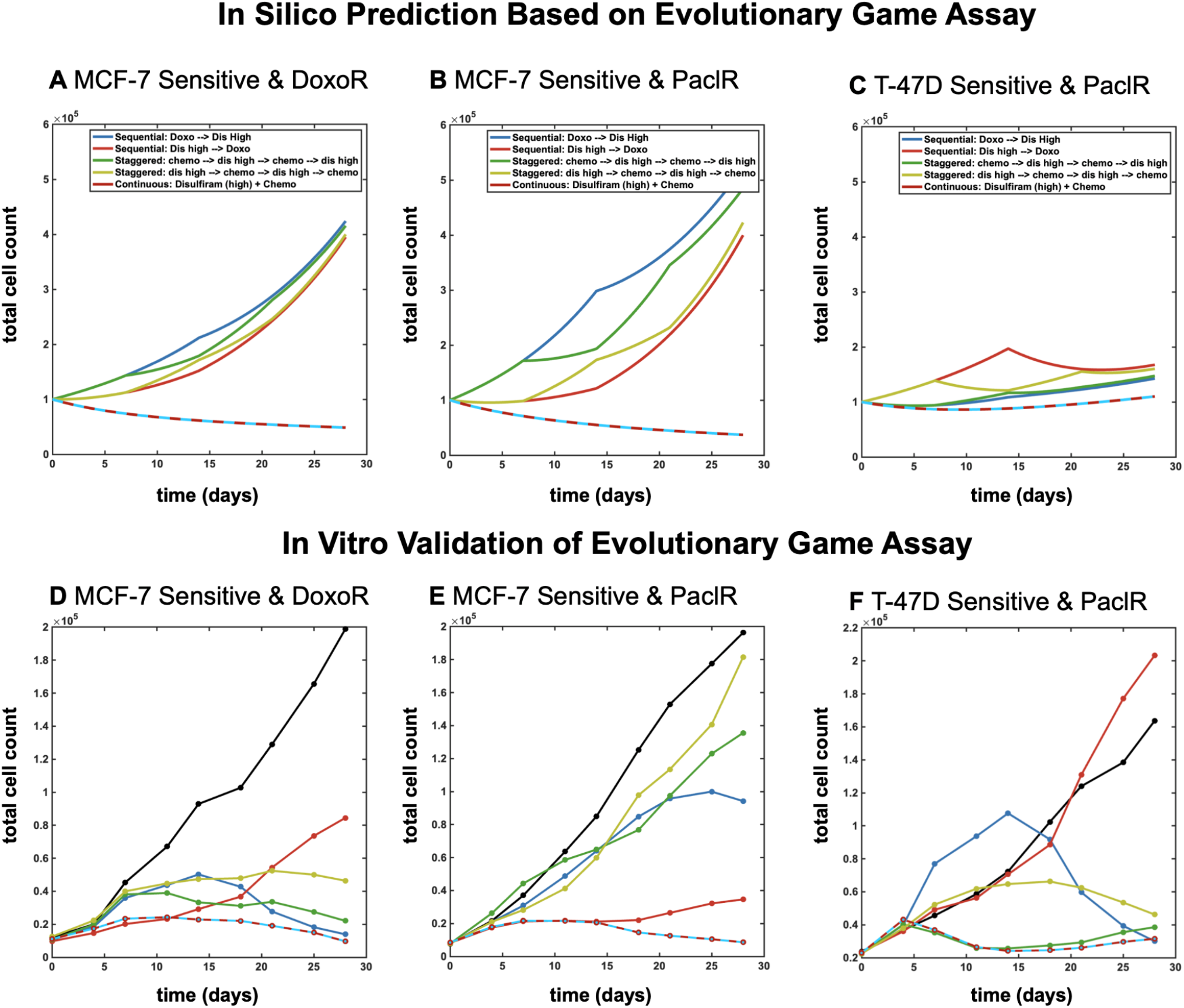
Model predictions for continuous and alternating treatments. Mathematical model simulations, parameterized in figure 4, shown for MCF-7 (**A, B**) and T47-D (**C**) cell lines, respectively. Top panels: *in silico* simulations show spheroid cell count over time for continuous mono- and combination therapy and alternating/sequential therapy for fast and slow switching. Bottom panels: *in vitro* validation of model predictions for spheroid cell count over time for all therapy regimens. Treatments include chemotherapy concentration of 25nM doxorubicin and 500nM disulfiram.

Theoretical results are experimentally validated in **Figure 5** (**Fig A, B, C**; bottom panel). Interestingly, in MCF7 lines, leading with disulfiram in alternating/sequential strategies produces more effective control of overall cell count compared with initiating treatment with chemotherapy. However, spheroid cell count is most negatively impacted under combination treatment out of all regimens. Therefore, the *in vitro* experiments confirm that combination therapy outperforms all sequential therapy regimens and is ideal across cell lines and initial sensitive-resistant fractions (shown left-to-right in bar groups).

## Discussion

A promising strategy for combating treatment resistance is to identify a candidate drug that can be classified as an evolutionary double-bind, whereby resistance to a first-line therapy confers sensitivity to a second-line treatment. In this work, we have identified disulfiram as the first reported example of an evolutionary double-bind with chemotherapy in ER+ breast cancer. To confirm the evolutionary double-bind, we developed an integrated mathematical-experimental framework to screen for promising candidate drugs.

Here, we developed resistant cell lineages over the course of long-term exposure to either paclitaxel or doxorubicin, and fluorescently labeled each cell line to track growth and treatment response over time. An extensive drug screen was performed using a range of candidate drugs either currently approved for use in breast cancer, exhibiting potential off-label use in oncological settings, or that inhibit pathways linked to resistance. From a panel of over 30 drug candidates, disulfiram was a leading drug found to be the most effective at inhibiting cell survival in all three chemo-resistant cell lineages compared to their respective chemo-sensitive cell lines. Cells resistant to paclitaxel harbored cross resistance to doxorubicin, but both resistant lines also harbored mutual sensitivity to disulfiram. This suggests a commonality between resistance mechanisms across these different chemotherapies. This commonality leads to a shared mechanism that disulfiram can exploit.

There are several important novelties implemented in this drug screening methodology. First, it is uncommon to quantify drug-synergy in both parental and evolved-resistance cell lines to note changes in collateral sensitivity after the onset of resistance. Secondly, it is an open question how drug-drug synergy relates to cell-cell interactions between parental and evolved-resistance lines. Finally, it is unclear how to best implement drug scheduling and timing for candidate evolutionary double-bind treatments. The integrated mathematical-experimental framework outlined above addresses all these key questions.

### Disulfiram as an evolutionary double-bind with chemotherapy

From our comprehensive drug screen, we identified disulfiram as a candidate drug to engage in an evolutionary double-bind, capitalizing on secondary vulnerabilities arising in chemo-resistant cells. We quantified synergy by performing dose-response analyses on both chemo-sensitive and resistant lines treated with disulfiram in combination with either paclitaxel or doxorubicin. The quantification of drug synergy (table 1) illustrates a moderate degree of heterogeneity across cell lines. Interestingly, synergistic potency is rarely seen in naive cell lines, but does appear in some resistant lines (e.g. MCF-7 DoxoR and T-47D DoxoR lines). Synergistic efficacy is more common and seen in both naive and resistant lines. Another promising result is that antagonism is rarely observed during disulfiram treatument for these lines. Interestingly, the presence of chemotherapy stabilizes the efficacy bifurcation of disulfiram observed when given alone. It is also important to note that even drugs without synergy may benefit from combination therapy due to an additive effect^47–49^.

The relationship between drug synergy in resistant lines and corresponding cell-cell interactions between parental and resistant lines is under-explored. To quantify cell-cell interactions, we performed the evolutionary game assay (EGA) to measure the frequency-dependent growth of parental and resistant cells when co-cultured and treated with disulfiram, chemotherapy, and combination therapy. The EGA analysis confirms that disulfiram inhibits the fitness (growth rate) of chemo-resistant lines, causing re-sensitization to chemotherapy (figure 4). This is true across both doxorubicin and paclitaxel. Combination chemo-disulfiram results in the lowest total tumor fitness, across all cell lines. These results suggest that the two drug combination is an effective combination with one drug to target sensitive cells (chemotherapy) and a second to target resistant cells (disulfiram). EGA results also suggest that the combination treatment would outperform any sequential treatment paradigm. This result was validated in vitro, confirming the optimality of combination treatment (figure 5).

Disulfiram is an FDA-approved anti-alcoholism drug reported to also have anti-tumor activity in several pre-clinical studies and clinical trials. Disulfiram has shown strong tumor-selective toxicity and inhibition of cancer cell growth^50^. Repurposing drugs approved for other indications and expanding them for use in oncological settings brings many advantages. Drug repurposing decreases significant time and costs traditionally invested in research and development. It also reduces the drug development time frame by providing valuable safety data demonstrated in previous clinical trials. Several clinical trials have investigated disulfiram as a combination therapy with chemotherapy in solid tumors. For example, there are active trials in metastatic pancreatic cancer (NCT02671890) and treatment-refractory sarcoma (NCT05210374), while a completed trial in recurrent glioblastoma (NCT02678975) showed no statistical significant difference in 6-month survival^51^. For these reasons, disulfiram presents as a promising drug in refractory metastatic breast cancer, prompting greater investigation for combination treatment with chemotherapy in our current study.

While any specific molecular mechanisms conferring vulnerability to disulfiram in resistant cells were not explored in the current study, future aims will center around elucidating mechanistic targets related to disulfiram in metastatic breast cancer. Disulfiram is known to suppress angiogenesis^52,53^, constrain tumor metastasis^54,55^, and modulate the immune microenvironment^56,57^. Previous studies have also shown disulfiram to be an inhibitor of PI3K signaling, often dysregulated in breast cancer, leading to suppression of cell proliferation and survival^58^. MDR drug efflux pump expression differences are often found in chemo-resistant cells compared to sensitive cells, and recent studies have also presented evidence that disulfiram may combat malignancy by inhibiting the ABC drug transport proteins involved in drug resistance^59–61^. Disulfiram has been shown to inhibit NF-kB (involved in epithelial-mesenchymal transition) and the self-renewal of breast cancer stem cells^62^. Disulfiram metabolites have also been shown to induce p53, a key player in maintaining the balance between self-renewal and differentiation, ultimately leading to apoptosis and cell death^63^. Therefore, while the exact acquired resistance trait targeted by disulfiram in these studies is not yet known, molecular aberrations in these pathways and processes may render these cells vulnerable to disulfiram.

To strengthen disulfiram as a candidate drug for potential application in clinical oncology settings, further *in vitro* as well as *in vivo* testing of combination therapy with disulfiram would be necessary to determine the optimal dosing, timing strategy, and toxicity concerns for future clinical trials. Combination disulfiram with chemotherapy resulted in the best drug response. Disulfiram alone was associated with a non-monotonic dose response (worse response for higher doses), which was not observed when combined with chemotherapy (figure 2).

Additional dosing schemes should also consider combination treatment of disulfiram along with alternative agents, given it is unlikely disulfiram would be clinically administered as a monotherapy. For example, previous studies have shown disulfiram-copper complexes can induce apoptosis in prostate cancer^64^.Through integration of experimental and mathematical models, this work highlights the importance of identifying targetable drug sensitivities in chemo-resistant settings and the potential to predict and avoid selection of resistant populations under therapy. Future investigations will explore further applications of the evolutionary game assay as it may be expanded to include other resistant cell lines, treatment strategies, and dosing concentrations. Elucidating the underlying molecular mechanisms involved in chemo-resistance may also allow for a deeper understanding of the mechanistic drivers of the observed evolutionary double bind and the potential for clinical applications of disulfiram in combination therapy.

## Methods

### Cell lines and reagents

The previously authenticated estrogen-receptor-positive (ER+) T-47D (ATCC, # HTB-133) and MCF-7 breast cancer cell lines (ATCC, # HTB-22) were maintained in RPMI+ 10% FBS+ 1% antibiotic–antimycotic or DMEM+ 10% FBS+ 1% antibiotic–antimycotic solution, respectively. The chemo-resistant cell line creation (paclitaxel-resistant T-47D; doxorubicin-resistant T-47D; paclitaxel-resistant MCF-7; doxorubicin-resistant MCF-7) was performed by culturing cells in regular maintenance media with weekly 24-hour exposure in chemotherapy-treated media, using doxorubicin (Selleck Chemicals, Cat. No: E2516) or paclitaxel (Selleck Chemicals, Cat. No: S1150), at different concentrations. Briefly, once a week and for a 24 hour window, cells were cultured in 70nM doxorubicin treated media (doxorubicin-resistant T-47D), 80nM doxorubicin treated media (doxorubicin-resistant MCF-7), 30nM paclitaxel (paclitaxel-resistant T-47D), or 80nM paclitaxel (paclitaxel-resistant MCF-7). This was performed over a 6-8 month course of time to develop resistance. Resistance against chemotherapy was detected by the alteration of the dose–response curve measured using CellTiterGlo Chemiluminescent Kit (Promega Corporation, Cat. No.: G7573). Cell lines were confirmed to be mycoplasma-negative using the Mycoalert PLUS Mycoplasma detection kit (Lonza, Cat. No.: LT07-703).

### Drug screening and dose response experiments with sensitive and resistant cell lines

T-47D and MCF-7 sensitive and doxorubucin or paclitaxel resistant cells were used in a drug screen and treated with various therapies. Therapies used in the drug screen are summarized in **Supp. Table 1**. 2,000 cells were plated in 384 flat bottom well plates (Corning, Cat. No. 142761) on Day 0 and treated 24 hours from plating (Day 1) for a duration of 72 hours. After 72 hours, collateral sensitivity against each drug was detected by measuring the viability of each line using CellTiterGlo Chemoluminescent Kit (Promega Corporation, Cat. No.: G7573). Relative viability of each resistant cell under each condition was normalized to parental sensitive cells under the same condition and were subjected to log2-transformation. All 2D dose response experiments following the drug screen for mono- and combo-therapy with disulfiram and chemotherapy were plated, drugged, and analyzed in the same manner and performed in triplicates.

### Lentiviral labeling of sensitive and resistant cells

Using lentiviruses incorporating distinct fluorescent proteins, we labeled T-47D and MCF-7 parental sensitive cells (venus; LeGO-V2) and chemotherapy resistant cells (mCherry; LeGO-C2). LeGO-V2 and LeGO-C2 vectors were provided by Boris Fehse (Addgene plasmids #27340 and #27339). Lentiviruses with fluorescent proteins were created using Lipofectamine 3000 reagent (Thermo Fisher Scientific) following manufacturer’s protocol. T-47D and MCF-7 sensitive and resistant cell lines were transduced with lentivirus using reverse transduction. Briefly, 1mL of polybrene-containing cell suspension of 200,000 cells were plated in a well of a 6-well plate. Previously, 0.5 mL of viral aliquot had been dispensed in plate. Following 48 hours of incubation at 37 °C with 5% CO2, cells were washed and given fresh regular culture medium. To select for fluorescence-activated cells, fluorescently labeled cells were flow-sorted after further subculture of transduced cells to attain homogenously labeled cell populations.

### Mono- and coculture 3D spheroid experiments

The 18-25 day experiments were initiated with fluorescently labeled sensitive and resistant cell lines in different compositions. For long-term T-47D and MCF-7 spheroid experiments, 5000 cells were plated in different proportions (100% sensitive, 75/25 sensitive-resistant, 50/50 sensitive-resistant, and 25/75 sensitive-resistant) in 96-well round-bottom ultra-low attachment spheroid microplate (Corning, Cat. No.: 4520). 24 h later, spheroids were washed and fresh medium including treatment drugs was applied (day 0). Spheroids were treated for a total of 18-25 days with imaging and media change performed every 4th and 7th day of the week. Spheroids were treated with either doxorubicin (Selleck Chemicals, Cat. No: E2516) or paclitaxel (Selleck Chemicals, Cat. No: S1150) at specified doses and various strategies (either continuous, staggered, layered, or sequential) as described in results and figures 3 - 5. Imaging was performed using Cytation 5 imager (Biotek Instruments) recording signal intensity from brightfield, YFP (for Venus fluorescence), and Texas Red (for mCherry fluorescence) channels. Raw data processing and image analysis were performed using Gen5 3.05 and 3.10 software (Biotek Instruments). Briefly, the stitching of 2 × 2 montage images and Z-projection of 6 layers using focus stacking was performed on raw images followed by spheroid area analysis. To quantify growth under these conditions, we measured fluorescence intensity or relative sensitive and resistant populations and growth of spheroid area over the total time of the experiment. For cell count calculations, a standard curve was created by measuring the fluorescence and area of spheroids, 24 hours after plating at different cell numbers (a range of 500 cells to 250,000 cells) and at different proportions (100% sensitive, 80/20 resistant-sensitive, 60/40 resistant-sensitive, 40/60 resistant-sensitive, 20/80 resistant-sensitive, 100% resistant). Further details regarding 3D cell count quantification can be found under **Cell number quantification** in Methods (below). All coculture experiments were performed in triplicates.

### Cell number quantification

The following method was used to estimate the number of cells of each type based on fluorescent imaging data. The number of cells, *N*_*i*_ of type i (*i* = [*S,R*]) is estimated by fluorescent intensity, *F*_*i*_, according to the following equation:

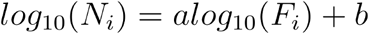

where a and b are constant values that differ for each cell line (MCF-7, T-47D) and for each evolved resistance line (parental, paclitaxel, doxorubicin). To correct for possible differences in per cell fluorescence, we regressed the fluorescence of pure cultures against the known numbers of S and R cells (Supp. Fig. S1), where the seeded cell number (x-axis) should match the predicted cell number (y-axis) across all initial sizes and sensitive-to-resistant ratios (circle color).

### Dose response

Treatment sensitivity is often modeled using a Hill equation. The Hill function can be derived by considering a weighted average of unaffected cells (E0), affected cells (E1) resulting at the equilibrium of a reversible transformation between these two populations, with corresponding dose-dependent rates of action^12^. The viability of a population of cells as a function of dose, x, can be written:

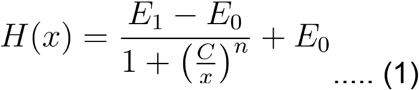

The function in eqn. 1 can be extended to consider two drugs with synergistic or antagonistic effects. Drug synergy or antagonism can be estimated by fitting a dose response surface, H(x_1_, x_2_) to a checkerboard dose response assay^10,11,65^. representing combinations of drug 1 and 2 at various concentrations, x_1_ and x_2_, respectively. For drugs that obey detailed balance, the two dimensional dose response is given by:

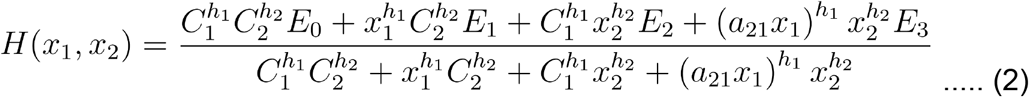

### Evolutionary Game Assay

The mathematical modeling follows the evolutionary game assay outlined in ref. ^25^, which we restate here briefly. Growth dynamics can be classified as having frequency-dependence if the fitness function changes with seeded ratio (frequency) of resistant cells. Fitness is a linear function between the monoculture growth rates for sensitive and resistant lines. The model has four parameters comprise the payoff matrix, **P**, providing a complete characterization of the fitness function of two competing cell lines. The evolutionary game assay is given by the following set of ordinary differential equations:

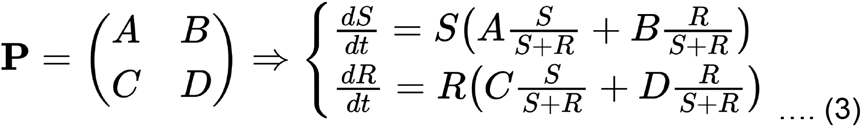

The two equations can be thought of as exponential growth with an instantaneous growth rate that is a linear function of the proportion of each cell type. For example, the equations above can be normalized to track the dynamics of the proportion of sensitive cells, which reduces to the replicator equation:

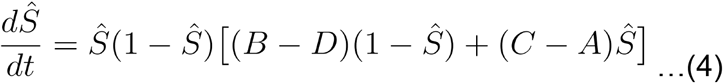

Thus, we define two important quantities that determine the selection dynamics in this model system: Δ_*S*_ and Δ_*R*_.

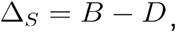

and

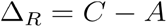

These two quantities represent the invasion fitness for each cell type when the proportion of that cell type is small. For example, if Δ_*R*_ > 0, resistant cells are expected to invade a population of sensitive cells. Whether that invasion continues to full dominance (resistant cells fully take over) depends on the value of Δ_*S*_. Finally, we denote the total tumor growth rate, *dS / dt* + *dR / dt*, (tumor fitness) at long-term equilibrium (i.e. *dŜ / dt* = 0), as *ϕ*\*.

## Acknowledgments

The authors gratefully acknowledge funding from the Cancer Systems Biology Consortium (CSBC) at the National Cancer Institute, U01CA232382, U54CA274507 and support from the Moffitt Cancer Center of Excellence for Evolutionary Therapy.

## Supplemental Information

**Figure S1:**
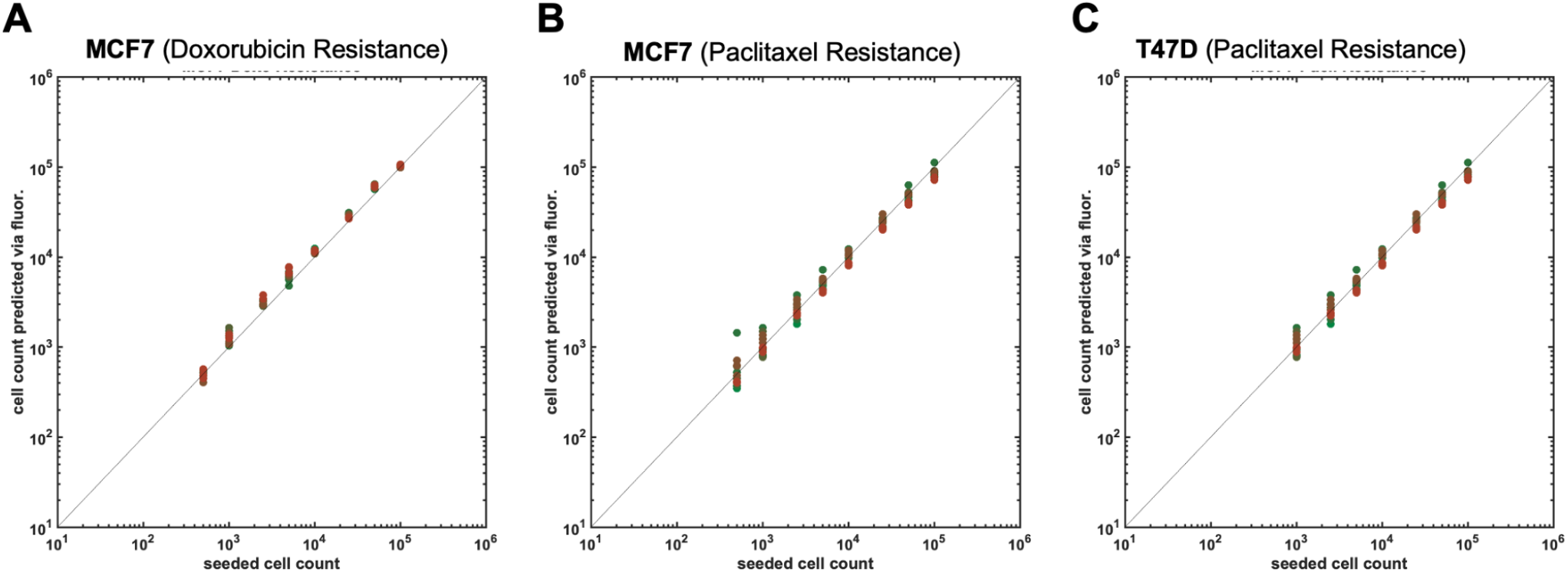
Validation of fluorescence intensity, F, to cell count, N, equation (see Methods). Known values of cells seeded (x-axis) are compared to predicted cell counts based on measured fluorescence intensity (y-axis). The model provides accurate cell count predictions, falling on the unity line (black line) across all initial sensitive-to-resistant ratios (color), repeated for (a) MCF7 DoxoR, (b) MCF7 PaclR, and (c) T47D PaclR lines.

